# Durable antibody responses in staff at two long-term care facilities, during and post SARS-CoV-2 outbreaks

**DOI:** 10.1101/2021.05.04.442699

**Authors:** Emily N. Gallichotte, Mary Nehring, Michael C. Young, Sierra Pugh, Nicole R. Sexton, Emily Fitzmeyer, Kendra M. Quicke, Megan Richardson, Kristy L. Pabilonia, Nicole Ehrhart, Bailey K. Fosdick, Sue VandeWoude, Gregory D. Ebel

**Affiliations:** Department of Microbiology, Immunology and Pathology, Colorado State University, USA; Department of Statistics, Colorado State University, USA; Columbine Health Systems Center for Healthy Aging and Department of Clinical Sciences, Colorado State University, USA

## Abstract

SARS-CoV-2 has had a disproportionate impact on non-hospital healthcare settings such as long-term care facilities (LTCFs). The communal nature of these facilities, paired with the high-risk profile of residents, has resulted in thousands of infections and deaths and a high case fatality rate. To detect pre-symptomatic infections and identify infected workers, we performed weekly surveillance testing of staff at two LTCFs which revealed a large outbreak at one of the sites. We collected serum from staff members throughout the study and evaluated it for binding and neutralization to measure seroprevalence, seroconversion, and type and functionality of antibodies. At the site with very few incident infections, we detected that over 40% of the staff had preexisting SARS-CoV-2 neutralizing antibodies, suggesting prior exposure. At the outbreak site, we saw rapid seroconversion following infection. Neutralizing antibody levels were stable for many weeks following infection, suggesting a durable, long-lived response. Receptor-binding domain antibodies and neutralizing antibodies were strongly correlated. The site with high seroprevalence among staff had two unique introductions of SARS-CoV-2 into the facility through seronegative infected staff during the period of study but these did not result in workplace spread or outbreaks. Together our results reveal that high seroprevalence rate among staff can contribute to herd immunity within a workplace and protect against subsequent infection and spread within a facility.

## Introduction

The emergence of SARS-CoV-2 and resultant COVID-19 pandemic threaten healthcare systems across the world [1, 2]. Long-term care facilities (LTCFs) are a significant venue for SARS-CoV-2 transmission and outbreaks and LTCF resident deaths account for almost half of all U.S. COVID-19 deaths to date [3, 4]. This is due to many factors including the communal nature of LTCFs and the high-risk health profile of residents [5, 6]. LTCF staff have the potential to introduce the virus into the facilities, where it can spread among staff, residents, and be exported back into the community. Additionally, staff at these facilities tend to resist vaccination [7-10]. We therefore began weekly SARS-CoV-2 surveillance testing of staff at LTCFs and observed significant facility-associated outbreaks [11]. In parallel with surveillance testing, we collected blood to determine seroprevalence, monitor seroconversion and characterize antibody responses in these populations.

Generation of specific, neutralizing and long-lived antibodies is a key component of adaptive immunity. Studies conducted after the SARS and MERS epidemics of 2003 and 2012 respectively, revealed that the majority of recovered individuals generated antibodies; however, it is unclear whether this immunity was sufficient to provide protection against re-infection [12, 13]. Many studies have sought to define the antibody response following SARS-CoV-2 infection [14, 15]. These include studies on hospitalized COVID-19 patients [16-19], asymptomatic individuals [20, 21], and retrospective serological studies [22-25]. The vast majority of infected individuals seroconvert and generate IgA, IgM and IgG-specific antibodies within 3 weeks of infection [15]. Age, sex, hospitalization, severity of infection, and other factors have all been shown to modulate the level, kinetics and durability of the antibody response following infection [21, 26-29]. Recent work has revealed that up to 7 months after infection, while absolute binding antibody levels might decline, neutralizing antibodies are long-lived, and persist at stable levels [30-36].

We therefore sought to characterize the antibody responses to SARS-CoV-2 in staff at two LTCFs by sampling serum at regular time intervals; during and post-outbreak. Using these samples, we measured antibody binding to two commonly used SARS-CoV-2 antigens, full length spike and receptor-binding domain (RBD), and neutralization of live SARS-CoV-2 virus. Our data clearly demonstrate the development of SARS-CoV-2 binding and neutralizing antibodies approximately 1-2 weeks post-infection, during the period of observation for the outbreak facility. Our data also reveal that the facility with high seroprevalence did not have any outbreaks during the study period, despite the introduction of the virus into the facility on two independent occasions. These results suggest that high seroprevalence (>40%) and levels of neutralizing antibodies can contribute to outbreak resistance through herd immunity. Additionally, we find that up to four months post-infection, neutralizing antibody levels are stable and durable.

## Materials and Methods

### Human specimens

This study was reviewed and approved by the Colorado State University IRB under protocol number 20-10057H. Participants were consented and enrolled in our study and promptly informed of all test results. Staff represent all job classifications, including those in direct patient care roles (nurses, physical therapists, etc.) and non-direct patient care roles (custodial, administrative, etc.).

### SARS-CoV-2 vRNA surveillance testing

Nasal swabs were collected, processed and tested for viral RNA as described previously [11]. Briefly, swabs were collected by trained personnel and placed in tubes containing viral transport media. RNA was extracted, and qRT-PCR performed using the CDC 2019-nCoV primers and probes [37], or the ThermoFisher Scientific TaqPath™ COVID-19 Combo Kit, under Food and Drug Administration (FDA) Emergency Use Authorization (EUA).

### Serum collection and processing

Whole blood was collected in BD Vacutainer® blood collection tubes (catalog #368660). Samples were incubated for 30-60 minutes at room temperature to ensure clot formation, spun at 1300xg for 10 minutes at 25°C with gradual acceleration and deceleration, sera were aliquoted and stored at −20°C. Prior to use, sera were heat inactivated at 56°C for 30 minutes, then stored at 4°C.

### Spike and RBD binding assays

RBD and spike ELISAs were modified from Amanat *et al*. [38]. Clear flat-bottom immune 96 well plates were coated at 2µg/mL with SARS-CoV-2 protein (Sino) and incubated overnight at 4°C. Samples were diluted 1:50 in diluent (1% milk powder, tween, PBS), and added to plates for 2 hr at room temperature after 1 hr of blocking (PBS, milk powder, tween). Positive controls included convalescent COVID-19 patient serum (gift of Raymond Goodrich) and monoclonal antibody CR3022 (Absolute Antibody). Charcoal inactivated pooled human serum collected in 2015 was used as a negative control (Jackson Immuno Research). Plates were washed 3X, then anti-human IgG HRP diluted 1:3000 (PBS, 1% milk, tween) was added for 1 hr. Plates were washed 3X, then indicator was added and incubated for 10 minutes (SigmaFast OPD, Sigma). Reactions were stopped with 3M HCl and plates were read at 490nm with a Multiskan® Spectrum spectrophotometer.

The cutoffs for classifying ELISA results as positive/negative were based on the average optical density (OD) values across two replicates. For each binding assay, the OD cutoff was specified as that which maximizes concordance with the SARS-CoV-2 neutralization assay results, specifically that maximizing the sum of the percent positive agreement (PPA) and the percent negative agreement (PNA), akin to Youden’s index. The resulting empirical PPA and PNA are 98% and 97% for the RBD binding assay, and 99% and 92% for the spike binding assay.

### SARS-CoV-2 neutralization assay

Vero cells were plated one day prior to infection. Heat inactivated sera were serially diluted in DMEM containing 1% FBS, mixed with ∼50 PFU SARS-CoV-2 (2019-nCoV/USA-WA1/2020 strain), and incubated for 1 hr at 37°C. Virus-antibody mixture was added to cells, incubated for 1 hr at 37°C, then overlaid with tragacanth media. Cells were incubated for 2 days at 37°C, then fixed and stained with 30% ethanol and 0.1% crystal violet. Plaques were counted manually.

### SARS-CoV-2 whole genome sequencing

Sequencing was performed as previously described [11]. Briefly, cDNA was generated using SuperScript IV, PCR amplification was performed with ARTIC tiled primers and Q5 High-Fidelity polymerase. PCR products were purified, libraries were prepared using KAPA HyperPrep Kit and unique index primers. Libraries were sequenced on the Illumina MiSeq V2 using 2 x 250 paired-end reads. Sequencing data were processed, quality checked, and consensus sequences determined.

## Results

### SARS-CoV-2 surveillance testing

We performed nasal surveillance testing for SARS-CoV-2 viral RNA of staff at two long-term care facilities over a 4-6 month period (Site A, July-Oct 2020, Site B, June-Dec, 2020, **Fig. 1**). Samples were collected at the workplace. Site A previously experienced a large outbreak in June immediately before our surveillance testing began, with 26 staff and 47 residents testing positive, whereas at Site B no symptomatic or asymptomatic cases had been diagnosed prior to our surveillance testing. Staff were tested at least once per week, approximately 180 unique individuals at each site participated in testing, with an average of 100 staff at site A and 85 staff at site B testing weekly (**Fig. 1a**). Positive tests and percent positivity varied by facility, with site A only experiencing two positive tests (from two different staff members) throughout their entire 17-week testing period (**Fig. 1b & c**). Site B experienced a large outbreak with over 15% of staff testing positive at its peak, and 24 unique staff testing positive throughout the 18-week study (**Fig. 1b & c**). We collected serum samples from staff at both sites every 3-5 weeks, spanning the 5-month surveillance period, including a timepoint immediately prior to and immediately following an outbreak in early September at site B (**Fig. 1c**).

**Figure 1.**
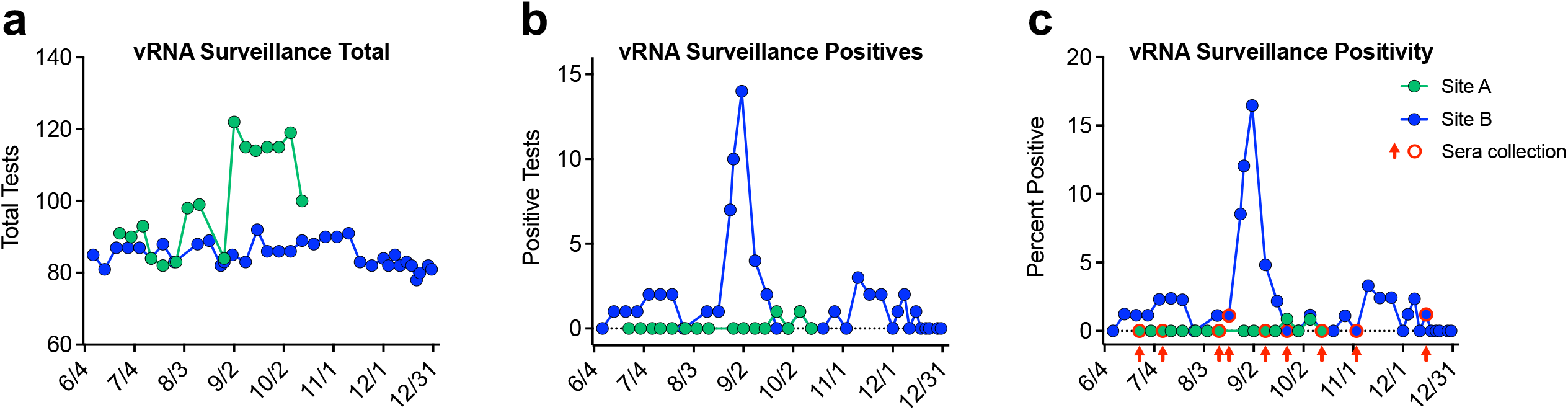
SARS-CoV-2 vRNA surveillance testing at two LTCFs. **a)** Total number of staff tested weekly as part of vRNA nasal surveillance testing. **b)** Number of positive vRNA tests recorded each week at sites A and B. **c)** vRNA positivity expressed as percent positive at each site. Timing of sera collections relative to surveillance testing are indicated by red circles and arrows.

### SARS-CoV-2 antibody binding and specificity

Sera from staff at both sites were evaluated for the presence and levels of polyclonal antibodies capable of binding to recombinant spike and receptor-binding-domain (RBD) proteins (**Fig. 2a**). At site A, spike and RBD binding seropositivity were approximately 40-50% over the 17-week study, with high agreement between the two antigens. Conversely, at site B, binding seropositivity at the start of the study, immediately prior to the large outbreak, was low (∼12%), but rapidly rose to ∼35%, post-outbreak (**Fig. 2a**). At site A, spike and RBD antibody binding levels gradually declined over the first 8 weeks, suggesting recent infection and progression from an acute to convalescent stage (**Fig. 2b**). At site B, binding levels quickly increased immediately following the outbreak and were stable over the following weeks (**Fig. 2b**).

**Figure 2.**
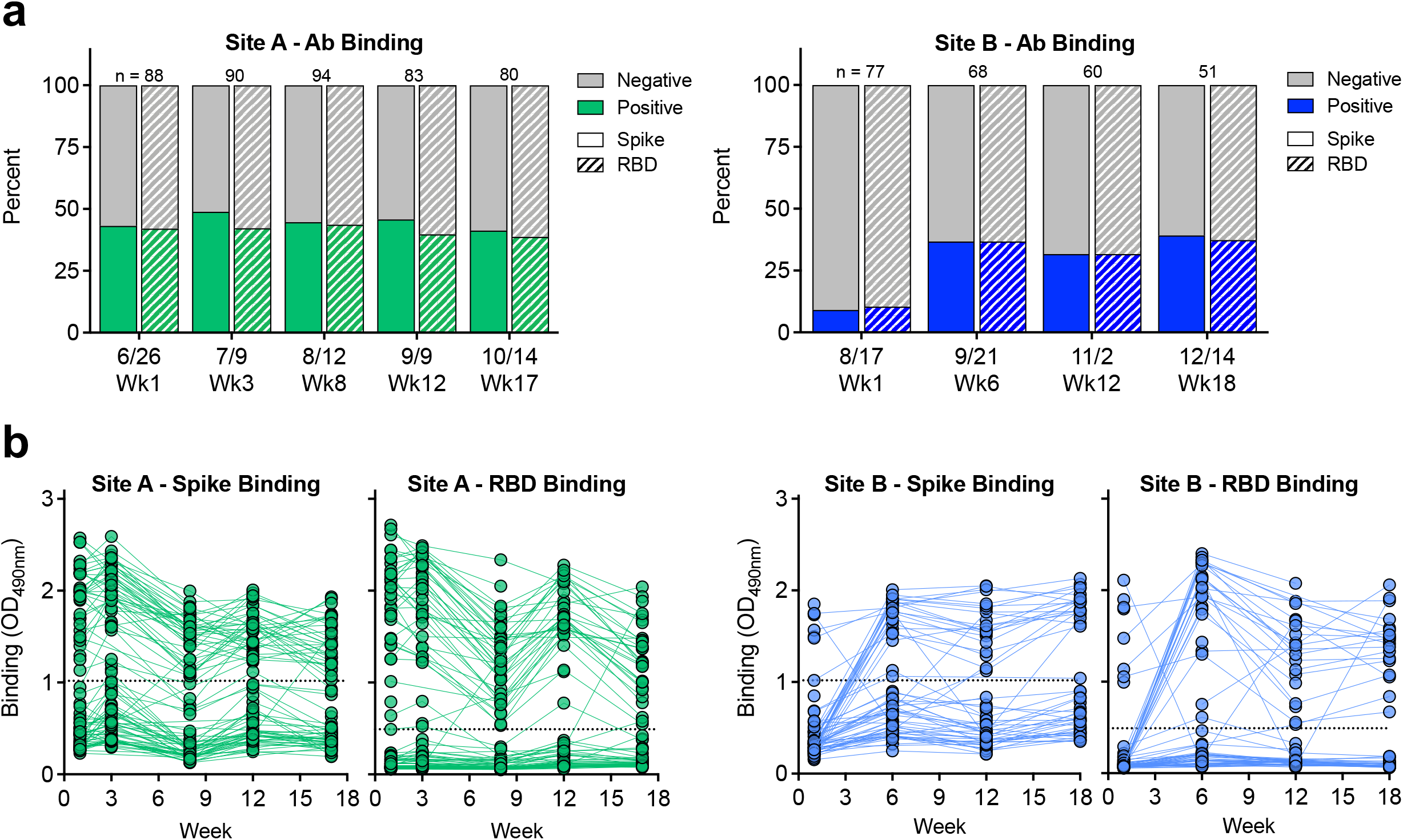
SARS-CoV-2 polyclonal antibodies bind spike and RBD. Polyclonal immune sera from sites A and B were evaluated for **a)** their ability to bind recombinant spike (solid) and RBD (dash) protein. N indicates number of samples tested each week. **b)** Level of spike and RBD binding as determined by absorbance reading. Dashed line represents Youden cut-offs.

### SARS-CoV-2 serum neutralization

Sera were next evaluated for their ability to neutralize live SARS-CoV-2 virus using a standard plaque reduction neutralization test, and their 50% neutralization titers were calculated (**Fig. 3**). In agreement with antibody binding results, site A had 40-50% neutralizing seropositivity which was maintained throughout the study, whereas at site B, rapidly increased from 10% to 35% between the first sample and subsequent weeks (**Fig. 3a**). At site A, neutralizing titers were highly stable over the 17-week study, whereas at site B, neutralizing titers rose as individuals became infected, decreased following the acute response, and were stable during convalescence (**Fig. 3b**).

**Figure 3.**
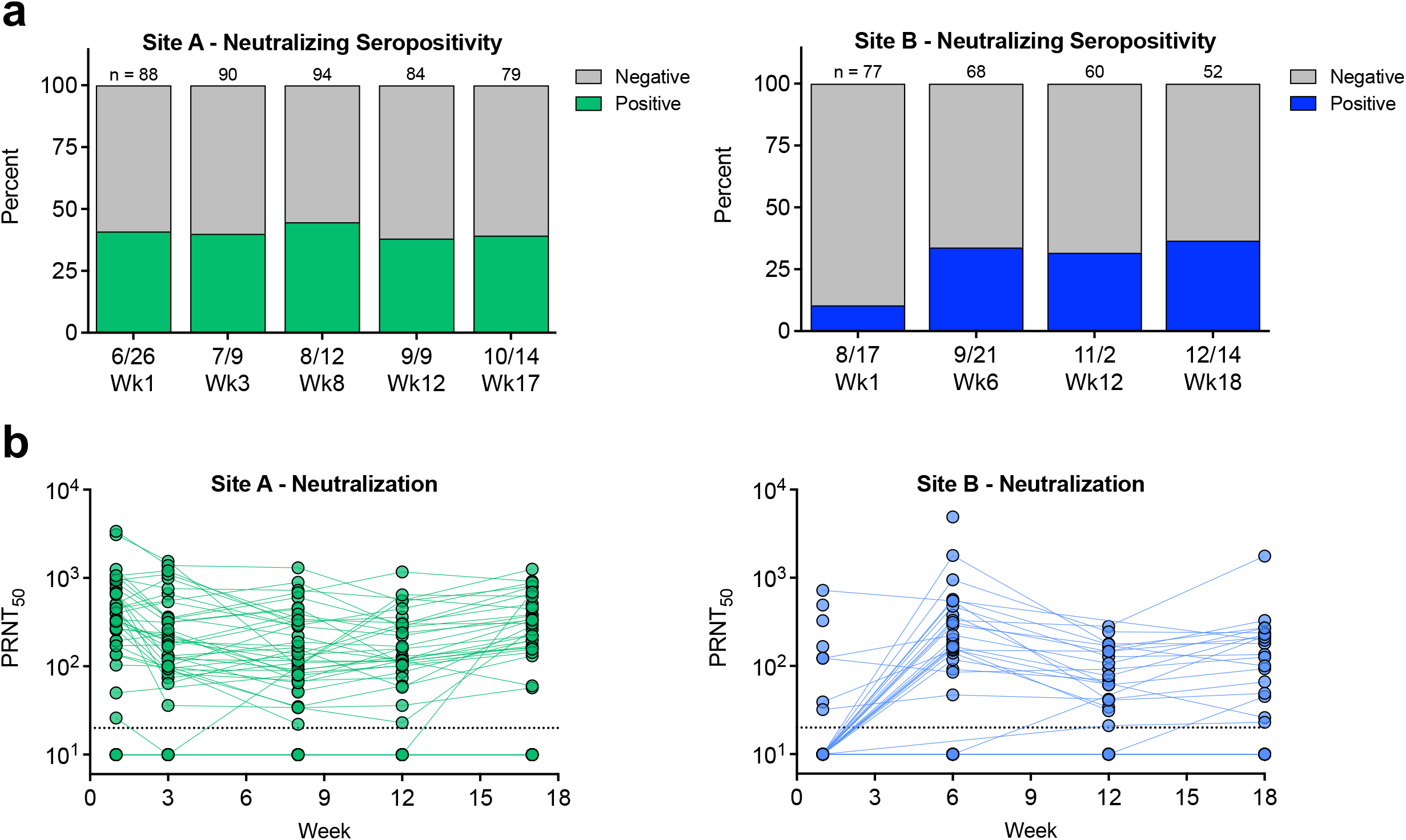
Polyclonal antibodies neutralize SARS-CoV-2 virus. Polyclonal immune sera from sites A and B were evaluated for: **a)** Ability to neutralize SARS-CoV-2 virus. N indicates number of samples tested each week. **b)** Neutralizing antibody levels over time. PRNT_50_ represents the serum dilution factor required to neutralize 50% of virus. Dashed line represents limit of detection (20). Non-neutralizing samples are graphed at half the limit of detection (10).

Neutralizing antibody levels of individuals at site B that were infected prior to the beginning of the study where highly stable over the 18-week study, suggesting they were infected weeks/months prior (**Fig. 3b**).

### Relationship between SARS-CoV-2 polyclonal antibody binding and neutralization

To better understand the relationship between binding and functionally neutralizing antibodies, spike and RBD binding levels and neutralizing titers were compared (**Fig. 4**). At both sites, spike and RBD levels were highly positively correlated (p<0.0001, Spearman r>0.7), suggesting the majority of spike antibodies bind within the RBD (**Fig. 4a & d**). At site A, there was a small population (3.9%) of samples with spike binding antibodies that are negative for RBD (**Fig. 4a**). Both spike and RBD antibody binding levels are highly correlated to neutralizing titers (p<0.0001, Spearman r>0.7) (**Fig. 4b, c, e, f**); however, at both sites, RBD-binding antibodies are more strongly correlated to neutralization (**Fig. 4c & f**).

**Figure 4.**
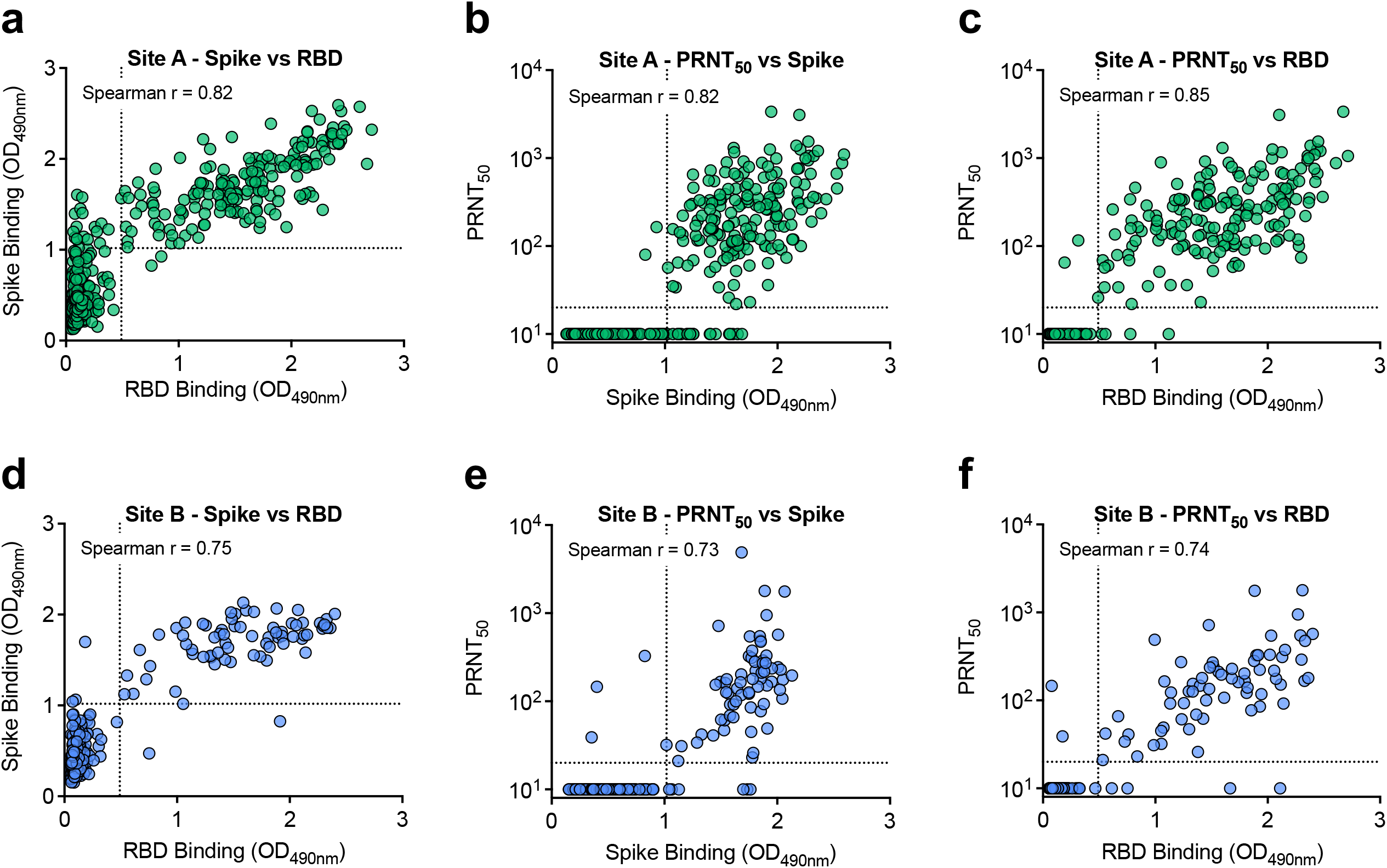
Spike binding, RBD binding and neutralizing antibody levels are highly correlated. Samples from site A (**a-c**) and site B (**d-f**) were graphed by spike and RBD binding levels (**a, d**), spike binding and neutralization titers (**b, e**) and RBD binding and neutralization titers (**c, f**). Spike and RBD dashed lines represent Youden cut-offs. PRNT_50_ represents the serum dilution factor required to neutralize 50% of virus. PRNT_50_ dashed line represents limit of detection (20). Non-neutralizing samples are graphed at half the limit of detection (10). Two-tailed, nonparametic Spearman correlation is noted in graphs.

### Kinetics of SARS-CoV-2 antibody levels post-infection

At site B, many individuals became infected and seroconverted during the course of the study. Therefore, in these individuals, we calculated the days post-infection (first positive vRNA nasal test) relative to seroconversion and levels of antibody binding and neutralization (**Fig. 5**). Spike binding antibody levels were high within 30 days of positive PCR test and remained high throughout the monitoring period (**Fig. 5a**). RBD binding levels were more variable and dynamic with some individuals generating RBD-specific antibodies within 10 days following infection, whereas one individual took over 60 days to seroconvert (**Fig. 5b**). Neutralizing antibody titers were also variable and dynamic across individuals, though most individuals generated high levels within a month following infection (**Fig. 5c**). In 85% of individuals, RBD-binding and neutralizing antibody levels decreased during the first 2-3 months following infection then stabilized (**Fig. 5b & c**, dashed lines). When comparing the relationship between binding and neutralizing antibodies stratified by timing post-infection, we again saw RBD and neutralizing antibody levels generally decrease ∼30 days post-infection, whereas spike antibodies were highly stable (**Fig. 5d**). Additionally, binding and neutralizing antibodies were highly correlated regardless of timing post-infection (p<0.0005).

**Figure 5.**
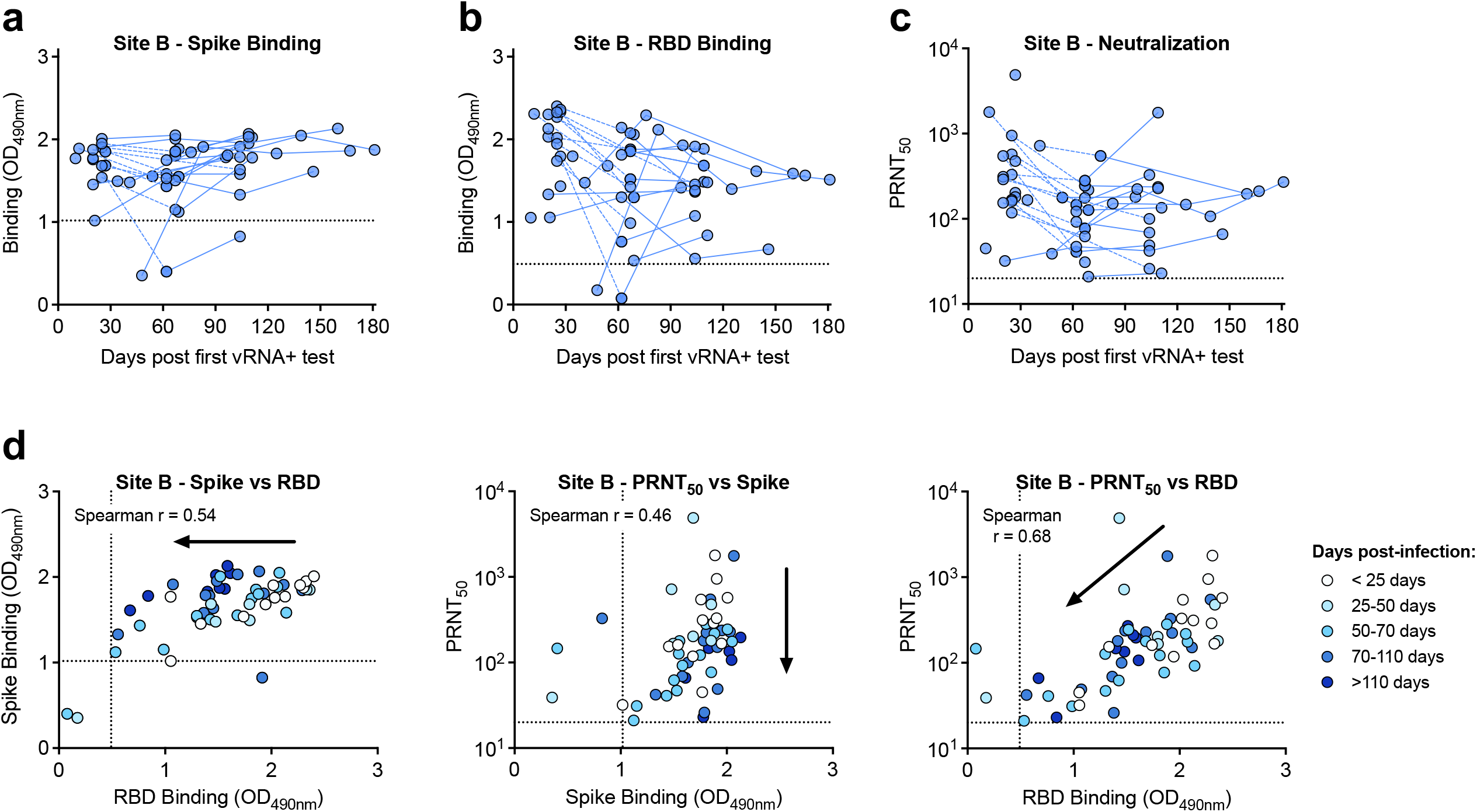
Trends in binding and neutralizing antibody levels vary over time. Individuals at site B who were infected during the course of the surveillance study were sampled up to 180 days post infection. **a**) Spike binding, **b)** RBD binding, and **c)** neutralizing antibody levels are graphed by days post-first vRNA positive test. **d)** Samples are stratified by days post-infection, and graphed by spike binding, RBD binding and neutralization titers. Arrows show trend of data over time. Spike and RBD dashed lines represent Youden cut-offs. PRNT_50_ represents the serum dilution factor required to neutralize 50% of virus. PRNT_50_ dashed line represents limit of detection (20). Non-neutralizing samples are graphed at half the limit of detection (10). Two-tailed, nonparametic Spearman correlation is noted in graphs.

### Phylogenetic analyses reveal lack of workplace SARS-CoV-2 spread

While site A did not experience any outbreaks during our surveillance testing, two individual staff members tested positive during the course of our study (**Fig. 1b**). These infections did not result in outbreaks or spread to other staff (**Fig. 2a & 3a**). The two individuals that tested positive for SARS-CoV-2 vRNA (two weeks apart on 9/22 and 10/6), provided serum samples in the weeks preceding their infections. Both individuals lacked detectable binding or neutralizing antibodies prior to infection and were thus immunologically naïve (**Fig. 6a & b**). To determine if the two viruses were genetically related, and therefore likely acquired from one another, viral genomes from the cases were sequenced. Both viruses contained shared single nucleotide polymorphisms (SNPs) relative to a reference strain (WA01), however they also contained 13 unique SNPs that strongly distinguish one from the other (**Fig. 6c**), suggesting two independent infections.

**Figure 6.**
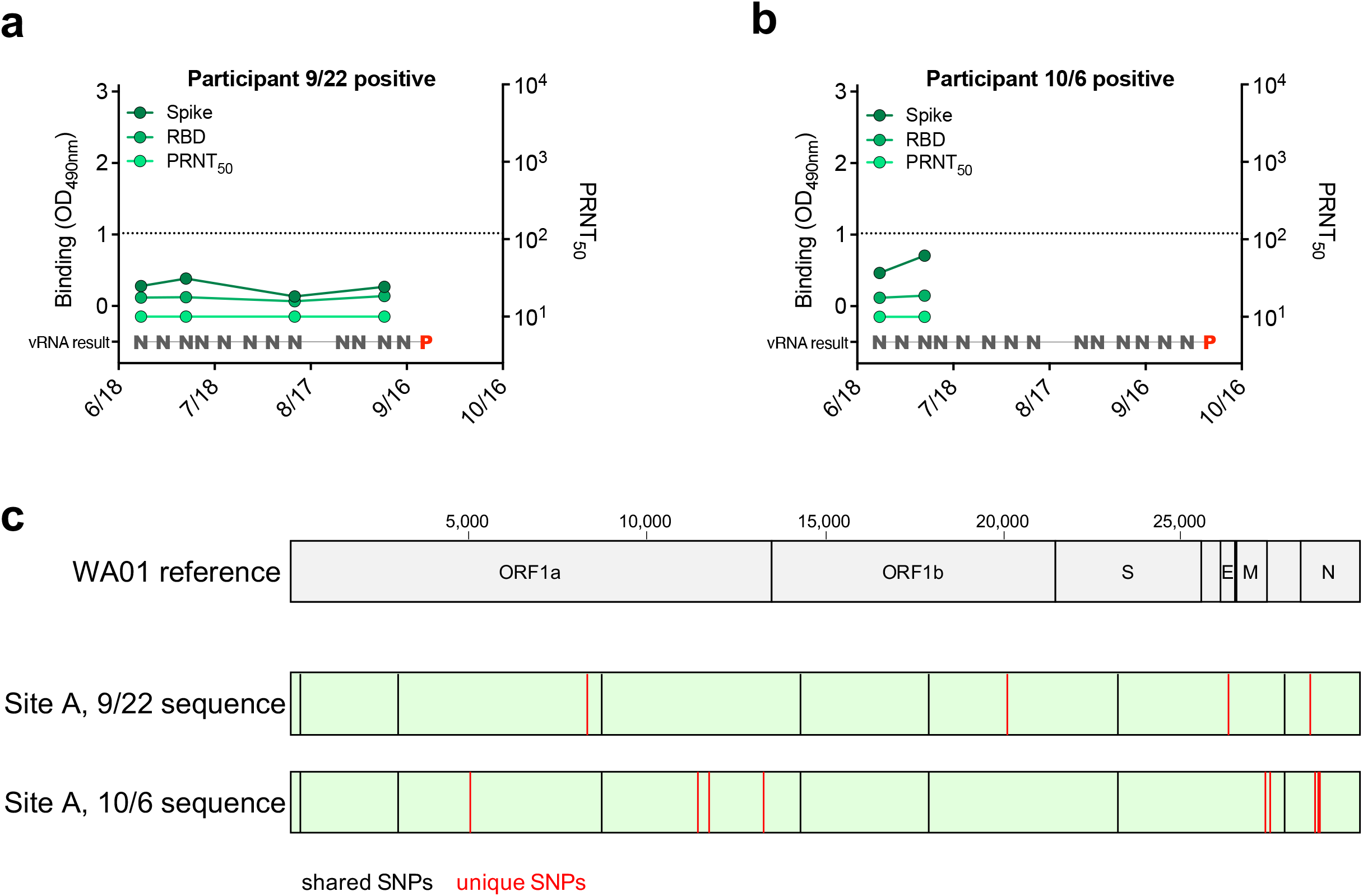
Two seronegative individuals at site A became vRNA positive with unique strains. **a, b)** Spike binding, RBD binding and neutralizing antibody levels relative to timing of surveillance vRNA testing indicated these two individuals were seronegative prior to infection (N – SARS-CoV-2 negative, P – SARS-CoV-2 positive). **c)** Viral RNA from positive surveillance testing was deep sequenced, and consensus sequence compared to the WA01 SARS-CoV-2 reference sequence. Single nucleotide polymorphisms (SNPs) shared between both site A sequences relative to reference are shown as black lines. Unique SNPs between site A sequences are shown as red lines.

## Discussion

Weekly surveillance testing revealed facility-specific SARS-CoV-2 infection rates. Site B experienced a large outbreak with 34 of the staff testing positive, whereas site A only had two positive tests out of greater than 1600 samples total. The high infection rate of staff at site B matches the incidence rates in staff at other LTCFs during outbreaks [11, 39-41], highlighting how quickly the virus can spread amongst long term care facility staff. Conversely, the low incidence of SARS-CoV-2 infection amongst staff at site A paired with the high seroprevalence suggests prior exposure and protection. Interestingly, at site A far more staff had antibodies than had previously tested positive for vRNA, suggesting a high fraction of asymptomatic infections, as has been documented in other facilities [40, 42, 43].

Two individuals at site A were vRNA positive for SARS-CoV-2 during the monitoring period. Full genome analysis of RNA recovered from these individuals revealed a significant number of genetic differences between the two isolates, suggesting they were acquired independently outside of work as two unique instances of community transmission. Our findings that staff at site A have high pre-existing seroprevalence (>40%) prior to intensive monitoring suggests this facility experienced a prior outbreak and had a level of herd immunity that limited spread of the virus from two positive staff members [44]. It is possible that other control measures and policies instituted at the time of monitoring, such as negative pressure isolation space [45], surveillance and monitoring systems and quarantine of positive staff [39, 46-48], environmental cleaning [49, 50], and others [51], additionally contributed to protection against outbreaks. It is notable that at both sites, seroprevalence reached a maximum of 40% during the study period, suggesting this might correspond to a level of herd immunity when coupled with other preventative measures.

Seroconversion and antibody levels were measured and characterized using three measures, binding to spike and RBD, and neutralization of live SARS-CoV-2 virus. We found that immediately following infection, antibody levels peaked during the acute phase then gradually decreased during convalescence. Neutralizing antibody levels were highly stable for at least 4 months post-infection, consistent with results reported by others [30, 34, 36]. Antibodies that bound to spike antigen were detected earlier and more consistently than antibodies binding to RBD; however, RBD-binding antibody levels correlated most strongly with neutralizing titers, a result reported in other studies [52-56]. Within our cohort, there are only four samples (0.57%) that neutralize SARS-CoV-2 but do not bind RBD. These likely neutralize through a mechanism other than blocking receptor interactions [57-59]. Since RBD-binding antibodies can be detected using high-throughput platforms such as ELISAs, whereas live-virus neutralization assays require BSL3 facilities, are lower throughput, and take longer, our observations that RBD-binding antibodies are strongly correlated with neutralization suggest the more convenient binding assay may, in some circumstances, serve as a substitute for functional anti-viral assays.

SARS-CoV-2 infection results in an immune response that includes the development of neutralizing antibodies [14, 15]. These antibodies provide some degree of protection against reinfection with SARS-CoV-2; however, their persistence and durability are unknown, and human correlates of antibody-based protection are lacking [60-62]. SARS-CoV-2 outbreaks at LTCFs can lead to high levels of seroprevalence that can limit spread within facilities [63]. Without complete herd immunity, there are still non-immune naïve individuals who can become infected and spread the virus, possibly leading to secondary outbreaks [64]. In our study, we observed that 40% seroprevalence in one facility, coupled with enhanced environmental controls, afforded apparent protection against subsequent outbreaks compared to a facility with low levels of pre-existing seroconverted workers. Due to the high risk of infection of vulnerable individuals, vaccination of staff and residents in LTCFs were among the highest priority for vaccination [65]. Immunity from natural infections, in addition to vaccine-elicited immunity has drastically reduced the burden of SARS-CoV-2 in many LTCFs [66-68]. This immunity paired with additional infection control measures will continue to reduce the incidence and prevalence of SARS-CoV-2 infection and mortality in these vulnerable facilities.

## Acknowledgements

This work was supported by the Boettcher Foundation and funds donated by the Colorado State University Colleges of Health and Human Sciences, Veterinary Medicine and Biomedical Sciences, Natural Sciences, and Walter Scott, Jr. College of Engineering, the Colorado State University Columbine Health Systems Center for Healthy Aging and the CSU One Health Institute. The authors also gratefully acknowledge the CSU Veterinary Diagnostic Laboratory for diagnostic support, and the participation of the workers in the facilities who participated in this study, without whom it could not have been completed.

